# Beyond redistribution: In-stream habitat restoration increases capacity for young-of-the-year Chinook salmon and steelhead in the Entiat River, WA

**DOI:** 10.1101/665604

**Authors:** Carlos M. Polivka, Shannon M. Claeson

**Affiliations:** Pacific Northwest Research Station, USDA Forest Service, 1133 N. Western Ave., Wenatchee WA 98801

## Abstract

We conducted snorkel surveys for juvenile salmonids in reaches of the Entiat River (Washington, USA) treated with engineered log jams (ELJs), and in reaches without treatments, to determine if habitat-unit-scale observations can identify whether restoration has increased the habitat capacity of a reach. The conceptual basis and field methodology emphasize fish density data (fish/habitat area in m^2^) from unrestored habitat within a reach treated with ELJs compared to surveys in 1) unrestored habitat in untreated reaches and 2) restored habitat in treated reaches. A Bayesian generalized linear model enabled us to quantify density differences among habitat types using advanced computational statistics. Modal density of young-of-the-year Chinook salmon (*Oncorhynchus tschawytscha*) and steelhead (*O. mykiss*) was at least 3.1-fold and 2.7-fold greater, respectively, in restored habitat compared with unrestored habitat for all treated reaches examined. To distinguish the density differences in those reaches as capacity increases rather than redistribution from poor habitat to good habitat, we compared density in unrestored habitat in both treated and untreated reaches. Here we found no differences for either species, confirming that the increased density in restored habitat units did not come from depletion of unrestored habitat in the same reach. We thus concluded that restoration increased the habitat capacity of the reach at the scale of pools created by ELJs.

---

In the United States, public agencies, academic researchers, and private consultants are engaged in activities related to the management and restoration of aquatic habitat to recover valuable fisheries (Katz et al. 2007; Roni et al. 2008), involving millions of dollars of public and other funds annually (Bernhardt et al. 2005; O’Neal et al. 2016). In the Pacific Northwest region, for example, managers have thus far implemented restoration projects in several hundred kilometers of stream habitat to improve rearing conditions for salmonid fishes. Use of large wood or other material to build in-stream habitat structures is one of the most common techniques to protect and recover salmon populations across the region (Roni et al. 2015; Hillman et al. 2016).

Large wood structures (engineered log jams; ELJs) are intended to promote processes that restore channel morphology (Davidson and Eaton 2013), increase pool availability (Roni et al. 2010), provide cover (Bond and Lake 2003a), and stimulate production of macroinvertebrate prey resources for fish (e.g., Hilderbrand et al. 1997; Kail et al. 2007). An important component of this restoration is the evaluation of its efficacy by demonstrating a biological response using the most robust metrics available (Block et al. 2001).

Habitat modeling tools such as Ecosystem Diagnosis and Treatment (EDT; Lichatowich et al. 1995; Lestelle and Mobrand 1996) and Physical Habitat Simulation System (PHABSIM; Bovee 1982) use fish-habitat correlations to guide restoration actions based on the assumption that specific habitat improvements will increase fish abundance and effectively increase fish populations. However, correlative analyses (e.g., multiple regression, ordination methods) may lead to overestimation of the effect that manipulation of habitat variables will have on fish abundance due to weak correlation coefficients, despite statistical significance (Shirvell 1989; Lammert and Allan 1999), complex interactions between habitat variables (Beechie and Sibley 1997), variable scales of inference (Feist et al. 2010), and model uncertainty (McElhaney et al. 2010). Furthermore, field surveys can show underutilization of “good” habitat, possibly due to responses to biological factors (e.g., escapement/seeding, food availability, predation risk, or competition) not measured as part of physical habitat surveys (Polivka 2005).

Thus, fish restoration effectiveness studies that rely upon the comparison of mean fish density in restored vs. unrestored habitats (e.g., Bond and Lake 2003b; Roni et al. 2008), often show small or no effects of restoration (Roni et al. 2008; Whiteway et al. 2010, Stranko et al. 2012; Hillman et al. 2016). When positive numerical responses are observed, the distributional patterns can vary greatly by species, year, and time period within a rearing season (Pess et al. 2012; Polivka et al. 2015).

Increased capacity to support fishes is one hypothesized outcome of restoration given the relatively consistent association between total habitat area and fish abundance (Hankin and Reeves 1988; Solazzi et al. 2000; Connor et al. 2001). Habitat capacity increases can be demonstrated by increases in total fish density (fish/m^2^ surface area, e.g.), relative to pre-restoration conditions, measurable at the scale of the restoration effort. Many post-restoration monitoring programs thus try to identify capacity increases via measurements of density increases in restored habitat. However, such observations may simply indicate that individuals have redistributed themselves within a reach from poorer habitat to restored habitat (Roni 2019). This may be attributable to the possibility of missing important spatial or temporal patterns in fish distribution among habitats with an observation design that is strongly linked to one scale or another (Fausch et al. 2002).

Restoration programs often aim to make effects detectable at a whole-reach scale (Roni et al. 2013; Roni et al. 2018a; Bennett et al. 2016); thus, many effectiveness studies are designed to compare fish density among restored and unrestored reaches. Although the reach scale is seen as a meaningful one for the measurement of many physical and biological processes (Frissell et al. 1986), it can be problematic because the effects of in-stream structures can sometimes be very localized (Polivka et al. 2019).

Tools such as the BACI (before-after-control-impact) design (Bernstein and Zalinski 1983) make it relatively straightforward to assess the effects of habitat restoration. The design offers the advantage that these effects can be distinguished from background variation common to all observations and from differences among reaches/sites (Popescu 2012). Challenges to using such a design include finding suitable reference sites and collecting comparable data between reference and treatment sites (Conner et al. 2016). Moreover, although BACI studies enable detection of effects at spatial scales from habitat units to whole reaches (Roni et al. 2018b), they are only possible when there are adequate pre-treatment data, which is difficult to ensure unless restoration projects are planned well in advance. At larger scales, such as comparing whole reaches, most effectiveness study designs, including BACI, are subject to replication issues (Stewart-Oaten et al. 1986; Conner et al. 2016), which can exacerbate difficulties in observing a density increase in restored habitat given high variability inherent in the study system (Underwood 1994). What is needed is a method that is scaled appropriately to detect effects at the scale of restoration, is robust to the variability in fish density that often confounds effectiveness study results, and that can identify real or apparent capacity increases, even when pre-treatment data are not available.

Studies at the patch scale address the replication of habitat units (e.g., pools, riffles, glides; Cramer and Ackerman 2009) in restoration treatments and can identify the extent to which the effects of restoration are localized (Polivka et al. 2019). Comparison of unrestored habitat patches in a treated reach against restored habitat patches (usually ELJ pools) in the same reach, and then against unrestored habitat patches in untreated reaches can indicate apparent capacity increases (Polivka et al. 2015). Furthermore, such a comparison can be made without pre-treatment data, but requires that other assumptions be met. Prior to treatment, stream reaches in which restoration is planned should have the same average fish density per habitat as the nearby, untreated reaches (Figure 1a), assuming that treatment and control reaches have similar geomorphic, hydrological and ecological characteristics. This assumption could be affected by seeding levels from spawning in the previous year (i.e., different numbers of juveniles resulting from year-to-year fluctuations in escapement), but can still be verified for untreated reaches sufficiently near the reach to be treated (Gowan and Fausch 1996; Kennedy et al. 2014). To identify a capacity increase when pre-treatment data are not available, post-treatment fish density in restored habitats is examined for one of these hypothetical patterns:

1. *Redistribution of fish within the treated reach*. Fish density is higher in restored habitats relative to unrestored habitats in a treated reach, but density is lower in unrestored habitats in the treated reach relative to all habitats in untreated reaches (which are, of course, also unrestored). This is due to the movement of fish to the restored habitats from unrestored habitats within the reach, such that total average reach density has not changed (Figure 1b).
2. *Type 1 capacity increase*. Fish density is higher in restored habitats relative to unrestored habitats in a treated reach, and remains equal across unrestored habitats in both treated and untreated reaches. In this case, the fish that occur in restored habitat add to the density observed in unrestored habitat and therefore represent an apparent capacity increase. (Figure 1c; Polivka et al. 2015).
3. *Type 2 capacity increase*. Fish density is higher in restored habitats relative to unrestored habitats in the treated reach, and higher in unrestored habitats in treated reaches relative to those in untreated reaches. In this case, restoration has increased capacity of *all* habitats in the treated reach (Figure 1d) and a density increase at the reach scale might be observed.

**Figure 1.**
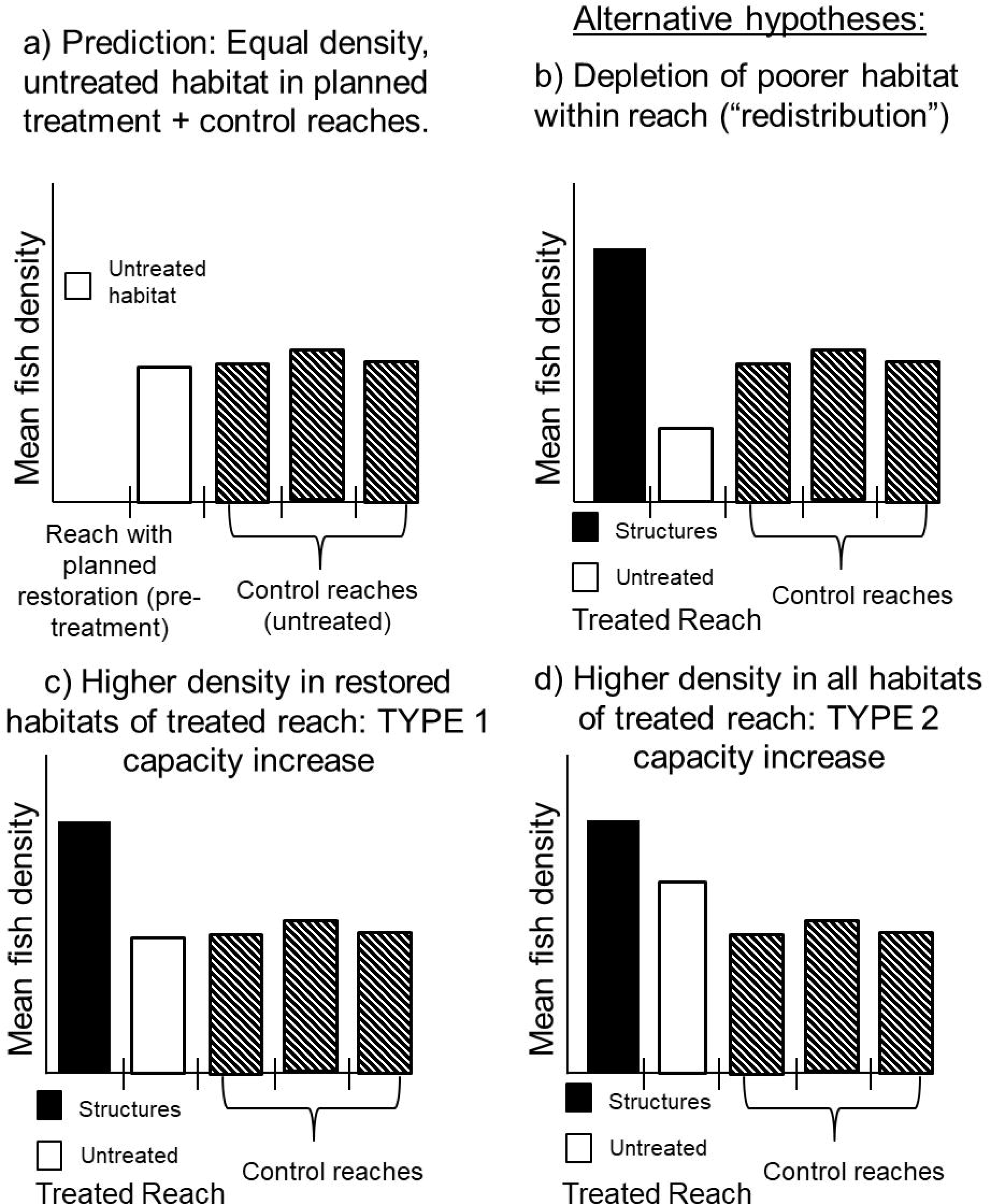
Theoretical fish densities at restored habitats (“Structures”) and unrestored habitats (“Untreated”) within a treated reach, relative to unrestored habitats within untreated reaches. A) Pre-treatment fish density at unrestored habitats in treated reaches and in untreated reaches. Post-treatment observations can involve: B) re-distribution of fish in the treated reach from unrestored habitat to restored habitat, C) additional fish in restored habitat, but no decrease in density in unrestored habitat relative to unrestored habitat in all untreated reaches, or D) increased density in restored habitat, and unrestored habitat in the treated reach is also increased relative to unrestored habitat in all untreated reaches.

Here, we apply the hypotheses in Figure 1 to post-treatment patterns of fish density from snorkel observations in eight reaches of the Entiat River, over multiple post-treatment years. We used a Bayesian analytical approach to deal with non-normally distributed density data – also a common issue in effectiveness studies. From this field methodology and analysis we were able to address whether restoration increases juvenile rearing capacity at both the habitat unit and reach scales in a manner distinguishable from redistribution of fish from poorer habitat to good habitat in a restored reach. Verification of this distinction can potentially aid study designs in systems where restoration effects are difficult to detect.

## [A]Methods

[C]*Study system*.—The Entiat River is a tributary of the Upper Columbia River in Washington State, USA (Figure 2). Its headwaters are in the Cascade Mountains and the river flows ∼ 69 river kilometers to its confluence with the Columbia River (49.6567 N, 120.2244 W; river km 778). Species targeted by restoration efforts include anadromous Chinook salmon *Oncorhynchus tschawytscha* and steelhead trout *O. mykiss*. The freshwater phase is an important component of the life cycle of these two species and in-stream habitat restoration is designed to enhance sub-yearling growth and survival.

**Figure 2.**
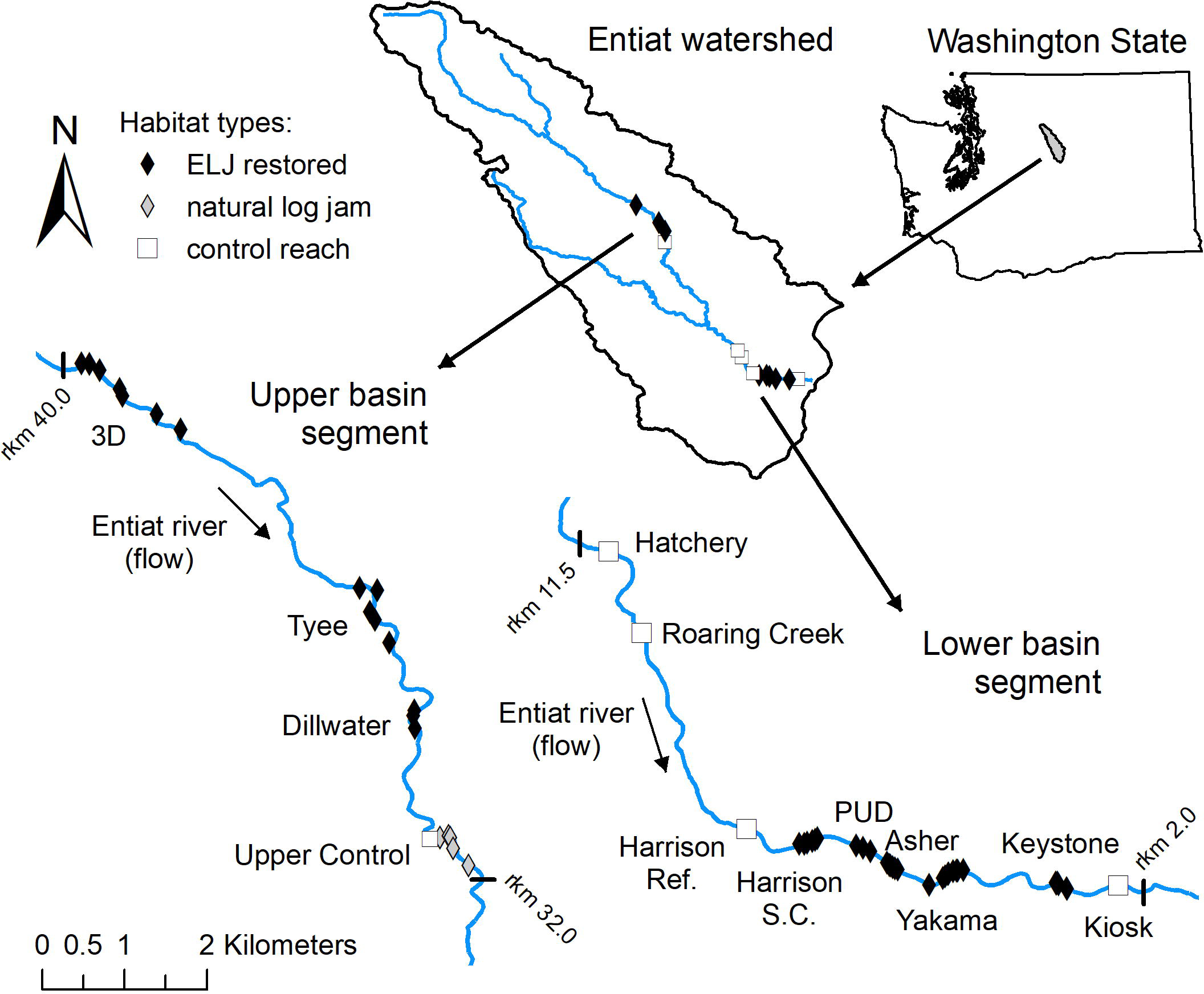
Locations of the treated reaches (with ELJs indicated by black diamonds) and untreated (control) reaches [natural log jams (Upper Control) or no wood at all (Hatchery, Roaring C., Harrison Ref., Kiosk)] in the upper and lower basin segment of the Entiat River. The upstream end of each control reach is indicated (white square). Additional details on reaches and treatments are given in Table 1.

**Table 1.** Lower and upper basin study reaches, their upstream reach location along the river, reach length, treatment type, and the number of wood structures (ELJ or natural) surveyed per reach. Unrestored habitat (N=15/reach) was also surveyed at each untreated and treated reach. Reach names were designated by restoration project sponsors and usually refer to a landmark or sponsoring entity.

| Basin | River | Reach | Reach name | Treatment (year) | Structures |
| --- | --- | --- | --- | --- | --- |
| segment | (km) | length (m) |  |  | (#) |
| Lower Entiat R. |  |  |  |  |  |
|  | 2.1 | 805 | Kiosk | Untreated |  |
|  | 2.7 | 322 | Keystone | Treated (ELJ 2014) | 6 |
|  | 4.3 | 322 | Yakama | Treated (ELJ 2014) | 8 |
|  | 5.1 | 644 | Asher/Milne | Treated (ELJ 2014) | 7 |
|  | 5.6 | 483 | PUD | Treated (ELJ 2014) | 3 |
|  | 6.4 | 322 | Harrison S.C. | Treated (ELJ 2014) | 6 |
|  | 7.2 | 644 | Harrison ref. | Untreated |  |
|  | 10.3 | 644 | Roaring Crk. | Untreated |  |
|  | 11.4 | 644 | Hatchery | Untreated |  |
| Upper Entiat R. |  |  |  |  |  |
|  | 32.8 | 483 | Upper Control (UC) | Untreated (natural) | 5 |
|  | 34.6 | 483 | Dillwater | Treated (ELJ 2012) | 3 |
|  | 36.2 | 805 | Tyee | Treated (ELJ 2012) | 5 |
|  | 39.9 | 1287 | 3D | Treated (ELJ 2012) | 7 |

Although some non-salmonid species and age 1+ steelhead are present in each of our study reaches, we focused our sampling and analysis on sub-yearling Chinook salmon and steelhead. The Chinook sub-yearlings in the Entiat are a mix of spring-run and summer-run and overlap during the study season. Genetic testing gives moderate to good probabilities of being able to distinguish runs (Degroseillier et al. 2018), but with some hybridization and spatial overlap, run type cannot be assumed. During the study years, total estimated spawning escapement of spring and summer runs (mean ± sd) in the previous season was 338 ± 142 and 258 ± 85 adults, respectively (Fraser et al. 2018).

There were two years in which Spring Chinook escapement and one year in which Summer Chinook escapement were slightly below 1 SD of the respective means. Thus our analysis did not consider annual differences in total seeding (e.g., density differences among habitats resulting from an unusually large Spring or Summer run) as a mechanism for habitat distribution of sub-yearlings reported below. Other salmonids are present in low abundance; cutthroat trout (*O. clarki*) and resident rainbow trout occur above the barrier to anadromy, which is upstream of our uppermost study reach. Bull trout (*Salvelinus confluentus*) move between the Entiat and mainstem Columbia and, although they can occur at our study sites, as can other resident freshwater fishes, none were observed during the study.

The restoration projects studied here were implemented in the Entiat watershed as part of a systematic program of restoration combined with post-treatment monitoring (Bennett et al. 2016). Treatment reaches (N =5) occurred in the lower part of the Entiat River basin, located between river km 2.7 and 6.4 (Table 1). Four untreated study reaches occurred among them with the most upstream reach at river km 11.4. The mean wetted width of lower basin reaches during the study was 23 m. Treated reaches in the upper basin (N = 3) occur between river km 34.6-39.9 with only one available control reach (“Upper Control,” UC; Table 1). The UC reach had naturally occurring log structures as well as untreated habitat and thus offered the opportunity to compare the effects on fish density of man-made ELJs with that of natural ELJs. The mean wetted width of upper basin reaches during the study was 21 m. Although the upper and lower basins are considered geomorphically distinct, (Godaire et al. 2009), both have similar substrate and gradient, and are used here only to identify the study reaches longitudinally.

[C]*Study reaches*—In the Entiat River, reaches were selected for treatment with ELJs by conservation agencies based on a combination of landowner willingness and recommendations following the completion of reach assessments for hydrologic conditions that were best suited for favorable outcomes of restoration (e.g., Sixta 2010). All reaches with ELJs were included in this study as treated reaches (Figure 2, Table 1). Untreated reaches were the nearest available reaches with public or private permissible access in which no habitat restoration structures were installed. We refer to treated and untreated reaches as such, and to restored and unrestored habitats as pools that were created/restored with ELJs versus river habitat units (not necessarily pools) that had no restoration structures.

The five lower basin reaches were treated with ELJs in July and August 2014, whereas the three upper basin reaches were treated in September 2012 (Table 1). ELJs in lower basin reaches created pools with a mean area of 38.0 m^2^ (range = 2.9-310 m^2^) and in the upper basin with mean area 79.1 m^2^ (range = 20.4 – 232.2 m^2^). Because fish activity and density in the Entiat River tend to be highest in July compared to other summer months (Polivka et al. 2015), post-treatment surveys in both upper and lower basins took place in July (lower basin 2015-2016; upper basin 2013-2016). The upper-basin reach “Tyee” not surveyed in 2016 due to a change in accessibility.

[C]*Sampling design*—Fish density data came from snorkel observations in reaches of the upper and lower basins designed after those in Polivka et al. (2015) where unrestored habitat units were selected in a spatially randomized manner in both treated and untreated reaches. Restored units (ELJ pools) are sampled in treated reaches as they were encountered during the sampling of unrestored units. The general approach was to establish a starting point at the upstream end of each reach regardless if treated or untreated. We selected 15 unrestored habitat patches (standardized to 15m^2^: 3 x 5 m) by first using randomly generated distance measurements following the method of Polivka et al. (2015). Random distances had a minimum of 10 m and a maximum distance of total reach length divided by 15 (e.g., “Kiosk” reach length of 805 m/15 = 54 m maximum longitudinal distance between unrestored habitat surveys). This ensured that all randomly-selected unrestored habitat patches occurred within and likely throughout the entire length of the designated reach. Surveys proceeded from the upstream end of the reach and each randomly generated distance was used to determine the distance downstream to the next survey patch from the previous. The survey took place on either the left or right river margin, selected via coin toss. Only habitat patches occurring on the stream margin were surveyed because mid-channel surveys have been found to provide little additional information in other salmonid streams (Beechie et al. 2005) and the river margin is where ELJs were placed. The standardized area of 15 m^2^ for unrestored habitat patches was often smaller than the pool area of ELJ-restored pools, but we account for the effect of area in the analysis (see below).

Patches typically consisted of pools or slower-flowing glides, but generally not riffles because riffles are not a substantial component of stream margin habitat in the Entiat River. Unrestored habitat units were surveyed by one snorkeler working in an upstream to downstream direction, visually identifying and enumerating all fishes present. Chinook and steelhead in the study reaches were typically age-0, but any age 1+ or larger steelhead were noted, when observed. In treated reaches, ELJ-restored pools were encountered during the movement downstream between unrestored habitat surveys. As ELJs were encountered, restored habitat surveys were conducted in the entire pool and total pool area was measured. For these, two snorkelers would survey the pool starting from opposite ends, because these pools tended to be larger and this would decrease the chance of fish being missed due to the increased habitat complexity of the ELJ. The survey crew would then continue downstream following the random distances to the next unrestored habitat survey. In the upper basin, the survey protocol was identical, with the exception that, in the control reach (UC), unrestored habitat surveys were conducted along with surveys at the naturally occurring log structures.

Turbidity does not affect detection of fish in this river as visibility for snorkeling is always >2–3 m at all but the highest flows, which permitted snorkelers to completely see the survey area. In unrestored habitats, lack of physical structure ensured accurate fish counts because the entire 15m^2^ area was completely visible (Polivka et al. 2015). However, the ELJ structures varied in their complexity and it is possible that fish could be obstructed from view, resulting in underestimation of density at restored habitats. This would make the estimate of any effect of restoration on density more conservative. [C]*Data analysis*—Data consisted of fish density (fish/m^2^) observations in pools created by or restored with ELJs, in randomly selected unrestored habitat in treated reaches, in randomly selected unrestored habitat in untreated reaches, and at natural structures in the UC reach in the upper basin. In each case, these were fish counts with accompanying measures of pool area. The key comparisons in our design (Figure 1) are 1) between restored and unrestored habitat in treated reaches 2) between unrestored habitat in both treated and untreated reaches and 3) in the upper basin between restored habitat in treated reaches and natural structures in the UC reach. To consider each species and upper and lower valley segments, this required a total of ten analyses: Comparisons 1 and 2 for two basin segments and for two species (8 analyses); plus comparison 3 for two species but for one basin segment (2 analyses). Because comparison 1 in the upper basin included pools created by natural structures and untreated habitat in the UC reach, comparison 3 identified whether there was a disproportionately stronger density response to natural structures vs. ELJs. We excluded age 1+ steelhead from the analysis as they were only ∼ 10% of total steelhead observed.

For each comparison, we fit a Bayesian generalized linear hierarchical model using the R software package BRMS (Bürkner 2017; R Core Team 2019; Stan Development Team 2019). Models all consisted of the same basic structure. Because we used count data by habitat unit and exploratory analysis indicated overdispersion of the data, due partly to the frequent observation of zero fish in unrestored habitat units, we assumed a negative binomial distribution and verified the modeled distribution using posterior predictive checks. Presence or absence of restoration structures (“Treatment”) was a fixed effect, but with an additional level in the upper Entiat accounting for the presence of natural structures in the “UC” reach. The interaction term year x stream reach was a random effect because each one is a grouping variable (2 years x 5 reaches, Lower Entiat; 4 years x 4 reaches, Upper Entiat). Pool area (log-transformed) was included in all models as an offset term given its positive relationship to fish abundance (Polivka et al. 2015) and to enable back transformation of model output values to density (fish/m^2^). We did not include a fixed effect term to look for differences among the treated reaches because pairwise comparisons involve reductions in the data that possibly lead to systematic lack of fit or, alternatively, over-report effect sizes (Head et al. 2015). During model specification, we found that the coefficients for year were close to zero. We did a Leave One Out (LOO) cross-validation without year and the model with year x reach term was more parsimonious with the data. This specification was also supported by \5^“’^ values and the convergence time for the different models.

We initiated the models with weakly informative priors (e.g., half student-*t* prior with 3 degrees of freedom and a scale parameter that depends on the standard deviation of the response after applying the link function, i.e., the defaults provided by BRMS) and for each we ran four chains for 2000 iterations, using the first 1000 iterations as a warm up and thinning by 1 iteration, giving a total of 4000 samples (1000 per chain). We plotted chains and parameter pairs for all models to make sure the chains converged and parameters were identifiable. We also examined model statistics to ensure that than 1.1._\5_“’ was less

To determine whether there were differences in fish density (D) across habitat types, we extracted the 4000 posterior estimates for the fixed effects (i.e., D_restored_, D_unrestored_, D_natural_), back-transformed the draws, and took the difference (D_restored_ - D_unrestored_). Using this vector of differences, we calculated a 95% highest (Posterior) density interval (HDI) using an algorithm described in Kruschke (2015). We considered fish density to be different among habitats if the 95% credible interval for that difference did not overlap zero. We used these same procedures to compare unrestored habitat in treated reaches with unrestored habitat in untreated reaches for each species and for the upper and lower basin segments separately.

## [A]Results

The two years of post-treatment sampling in the lower basin yielded a total of 328 surveys, 58 of which were at the ELJ-restored pools distributed among the five treated reaches, and 270 of which were in randomly-selected unrestored habitat units in treated and untreated reaches, combined. The four years of post-treatment surveys in the upper basin yielded 294 surveys, consisting of 52 at the ELJ-restored pools and 17 at the naturally occurring log structures in the UC reach. The number of structures in that reach varied for each year of the study due to wood aggregation and wash-out (N = 3 (2013), 5 (2014, 2015), and 4 (2016)). Across all study reaches and habitat units, we observed 1504 sub-yearling Chinook and 887 steelhead in the lower basin and 1189 sub-yearling Chinook and 273 steelhead in the upper basin. Overdispersion of the data was exacerbated by surveys with zero fish (no Chinook in 145 and 221 surveys, and no steelhead in 148 and 236 surveys, in the lower and upper basins, respectively) and by positive counts varying across orders of magnitude.

Pools created by restoration with ELJs showed higher fish density than unrestored habitat patches. In the lower basin segment, restored habitat in the five treated reaches had, on average, 2.9-fold greater density of sub-yearling Chinook salmon and 1.7-fold greater density of steelhead than unrestored habitat in those same reaches (Figure 3A).

**Figure 3.**
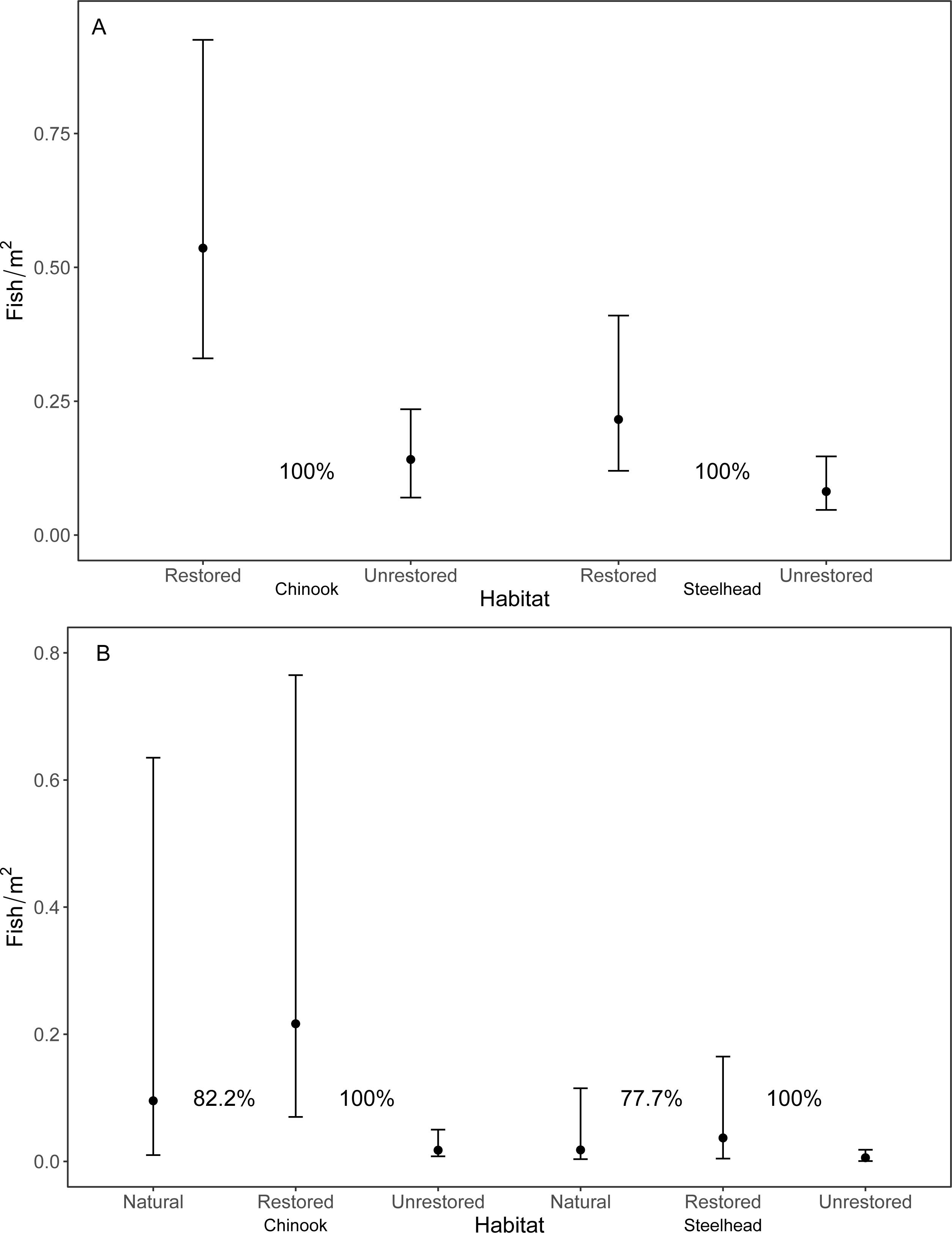
Estimate of modal Chinook and steelhead density (± 95% confidence interval) from a generalized linear model of fish counts from A) five reaches treated with restoration structures in the lower basin of the Entiat River and B) three treated reaches in the upper basin. Comparisons are between habitats restored with ELJs and unrestored habitats in the same reach. In the upper basin, an additional comparison was made between modal density in habitats restored with ELJs and habitats created by natural wood structures. Displayed are the percentages of draws from the posterior distribution in each habitat type in which the difference in fish density between habitats (ΔD) was > 0 (see Table 2 for details).

**Table 2.**
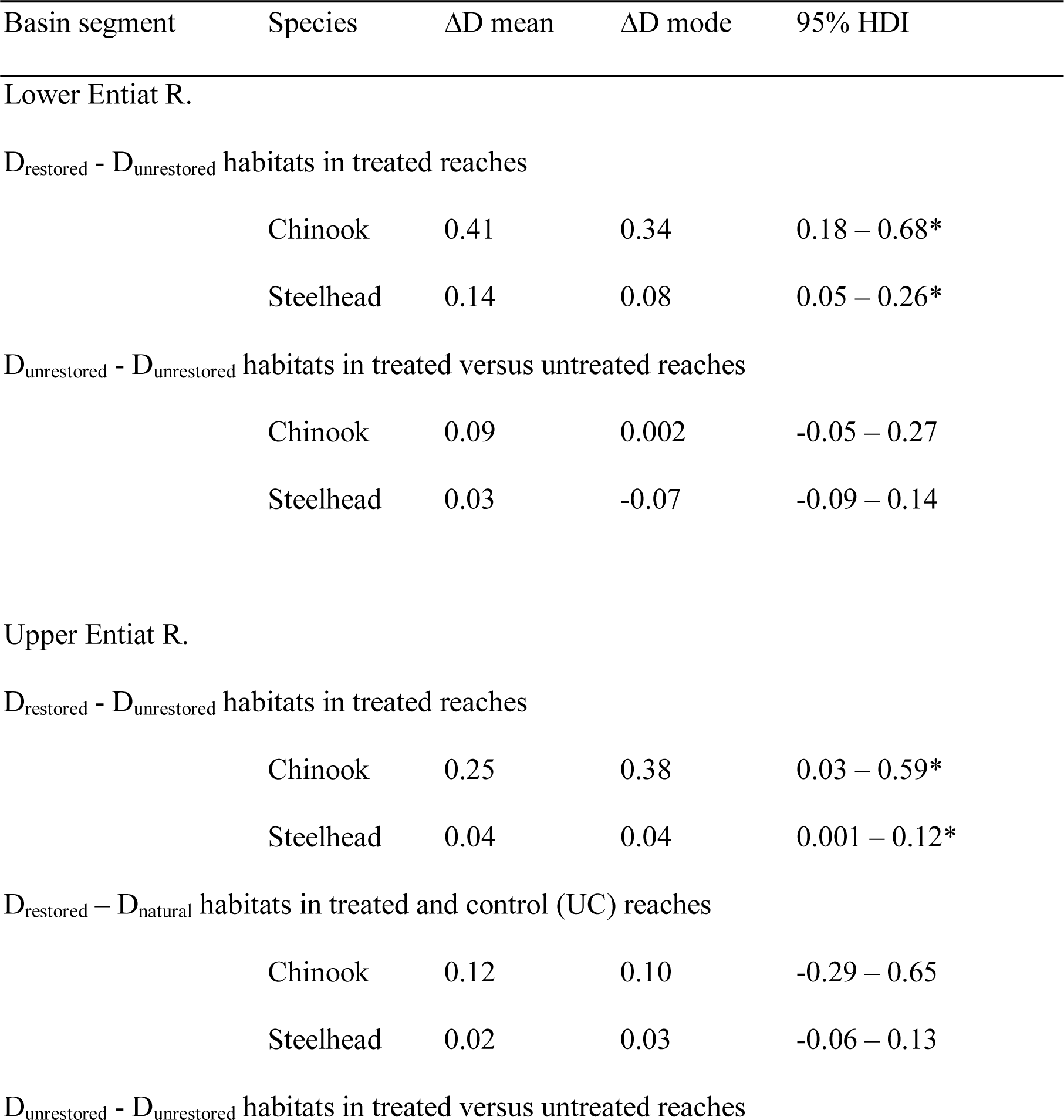
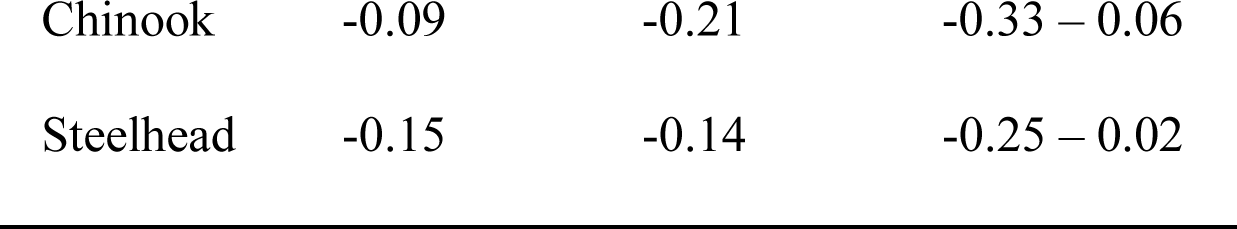
Mean and modal differences in fish density (D) drawn from the posterior distribution of generalized linear models fit to survey data in each indicated habitat type from treated and untreated reaches in the Entiat River. The 95% highest density interval (HDI) of each comparison is given. HDIs that do not overlap zero (*) indicate meaningful differences in density (ΔD).

Draws from the posterior distributions of density in each habitat (D_restored_, D_unrestored_) led to a distribution of density differences between habitat types (D_restored_ - D_unrestored_) with a 95% credible interval > 0 in 100% of the draws for both species (Table 2). In treated reaches of the upper basin, restored habitat had a 13-fold higher density of Chinook and a 6.8-fold higher density of steelhead than unrestored habitat in the same reaches (Figure 3B). Despite the magnitude of those differences in overall means, the draws from posterior distributions led to a 95% credible interval that nearly overlapped zero in both species (Table 2). There were no density differences for either species between restored habitats in treated reaches and habitats with natural structures in the UC reach (Figure 3B), indicating that ELJ structures are good analogs for natural structures, at least in this study system. This also indicates that natural structures did not meaningfully contribute to the observed overall difference between restored and unrestored habitat in the upper basin.

For the second key comparison in detection of a capacity increase, unrestored habitat in treated reaches did not differ in density (of either species) from unrestored habitat in untreated reaches in the lower basin (Figure 4A), nor were there density differences between unrestored habitat in treated reaches in the upper basin segment and unrestored habitat in the UC reach that only had natural structures (Figure 4B). In both the upper and lower basins, the lack of a difference between unrestored habitats in treated reaches and in untreated reaches indicates a Type 1 capacity increase (Figure 1). In the lower basin, however, Chinook density was higher in unrestored habitats of treated reaches than in unrestored habitats of untreated reaches in nearly 93% of the draws from the posterior distribution. Thus lower basin restoration could be approaching a Type 2 capacity increase that may become evident in later post-restoration years. From both pairs of figures it is clear that the lower basin has a higher density of both species than the upper basin.

**Figure 4.**
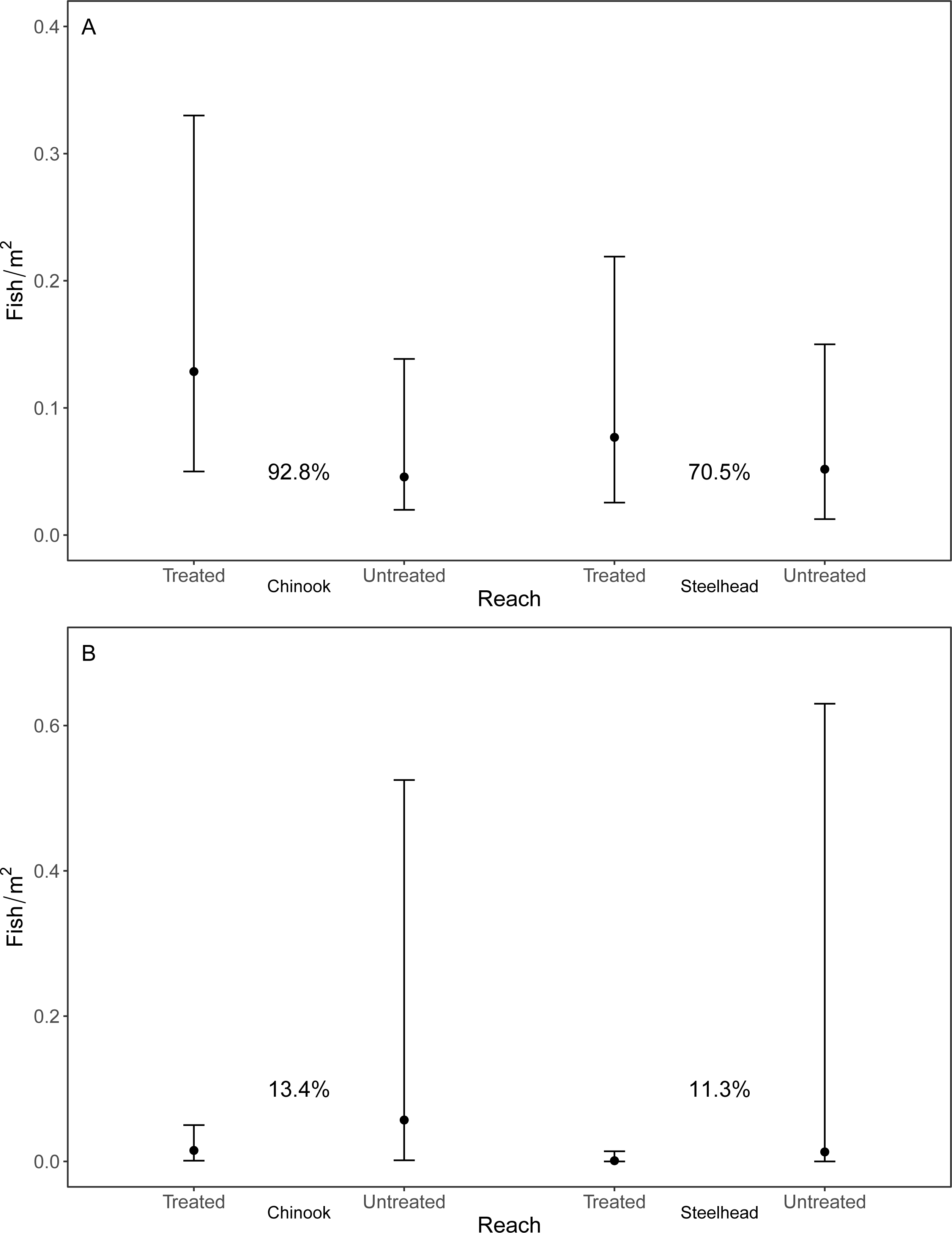
Estimate of modal Chinook and steelhead density (± 95% confidence interval) from a generalized linear model of fish counts comparing unrestored habitat in A) five each of treated and untreated reaches in the lower basin of the Entiat River and B) three treated reaches and one untreated reach (UC) in the upper basin where habitats with natural log structures occurred. Displayed are the percentages of draws from the posterior distribution in each habitat type in which the difference in fish density between habitats (ΔD) was > 0 (see Table 2 for details).

## [A]Discussion

Although it is relatively easy to show that fish are attracted to in-stream habitat restoration structures, there are challenges to demonstrating that increased fish density represents an increase in habitat capacity. The empirical basis of the method presented here serves effectively when pre-treatment data are lacking on fish density in the area to be restored, thus making BACI designs intractable. Like BACI, however, we relied upon an assumption that is difficult to demonstrate conclusively. Nevertheless, in this case we had a sufficient number of treated and untreated reaches to distinguish between a capacity increase and simple redistribution of fish. Moreover, we provided an analytical framework that addresses the issue of overdispersion in data that occurs, for example, when there is a considerable amount of unused habitat resulting in observations of zero fish. The field methodology involves sampling at the habitat unit (“patch”) scale (Frissell et al. 1986), which may be difficult to extrapolate to larger scales as may be desired by restoration planners (Roni et al. 2008). Our method is robust to many of these concerns and even showed a trend toward a larger scale effect in one group of treated reaches in our study system. Finally this study design is broadly applicable to myriad restoration projects that lack rigorous pre-treatment data, and combined with pre-treatment data would be even more powerful.

Our approach rests upon the assumption that unrestored habitat in treated and untreated reaches show the same value of the response variable (fish density or presence/absence). A BACI design does not depend on this assumption and only requires that a treated area show a greater positive change, relative to an untreated area, following the treatment (Roni et al. 2013; Conner et al. 2016; Roni et al. 2018ab). However, stream restoration is often opportunistic as it is dependent upon the convergence of habitat in need of improvement with feasibility of implementing restoration treatments; thus, pre-treatment data are not always available. However, our key assumption could be affected by differences among reaches prior to treatment. For example, if the treated reach had lower density than other reference reaches prior to treatment, then a density difference between restored and unrestored habitat within that reach could be interpreted as redistribution, thus underestimating the positive response to restoration. Given that we surveyed multiple untreated reaches in the lower basin and found that unrestored habitat there did not differ from unrestored habitat in treated reaches, it is unlikely that the extent to which we observed a density increase in restored habitat was affected by unobserved variation.

We show that what happens in unrestored habitat gives information about how to interpret observed density increases in restored habitat. However it also depends on the availability of multiple suitable control reaches, which may not be available under some restoration programs or in some watersheds. In our case, land ownership and local physical conditions resulted in only a single reference reach being available in the upper basin, further complicated by the presence of natural log structures. Thus we couldn’t be certain of a Type 1 vs. Type 2 capacity increase in the treated reaches in the upper basin because we were not able to sample reaches without structures, man-made or natural. This reference reach did, however, provide the advantage of showing that the ELJs are, at least, good analogs for natural structures.

The mobility of salmonids and the dependence of spatial scale on their relationships with habitat characteristics (Feist et al. 2003, 2010), mean that movement and habitat settlement patterns may affect comparisons of unrestored habitat among treated and untreated reaches. Ideal free distribution theory (Fretwell and Lucas 1969; Morris 1988) describes how habitat quality affects settlement of habitats at varying densities. Depending on whether the difference among habitats is qualitative or quantitative, the density difference in individuals between higher quality habitats and lower quality habitats may be larger at low total population density than when population density is higher (Morris 1988). In this study system, years with low total fish density could therefore result in large density differences between restored and unrestored habitat in treated reaches if restored habitat is settled preferentially (quantitative difference; Morris 1988). Moreover, all unrestored habitats in untreated reaches would be settled at about the same rate, and comparison of treated and untreated reaches might thus show that unrestored habitat in treated reaches had lower density than in untreated reaches. This would give the appearance that the density difference between restored and unrestored habitat in a treated reach was simply due to redistribution, leading to underestimation of the habitat improvement by restoration. Variation in the total number of sub-yearling Chinook seeking habitat can result from year-to-year differences in escapement. These differences were not substantial in this system during the study years, but may need to be considered elsewhere.

Statistical issues in both this and with BACI designs may result from low replication (Conner et al. 2016). This was particularly an issue at the Dillwater reach in the upper basin where few structures were installed to begin with, some of which were above the water line during low late-summer flows. This supports the need for analytical techniques, such as ours, capable of combining multiple treated reaches and years to determine restoration efficacy. We frequently observed zero fish at the habitat scale; however, the Bayesian GLM fitting routine precluded the need for zero-inflated frequentist models (e.g., Zuur et al. 2009). Moreover, it afforded a robust method of back-calculating fish density and examining the entire posterior distribution of the density estimate for meaningful differences in density among habitat types. Bayesian approaches can also improve the interpretability of results from BACI designs while dealing with replication issues (Conner et al. 2016).

Despite apparent improvement of habitat capacity observed here and in Polivka et al. (2015), a given density increase associated with ELJ restoration can be difficult to observe at the reach scale. In a focused study at three of the five lower basin reaches studied here, Polivka et al. (2019) demonstrated that observed density increases at ELJ-restored pools are very localized, sometimes indistinguishable only 5-10 m away, or by increasing the sampling area around a restored pool to only 2X the area of the pool. This does not necessarily negate the apparent capacity increases observed here; it merely demonstrates that there can be a large difference between restored and unrestored habitat in a treated reach. Monitoring programs often emphasize a reach-scale perspective based on earlier recommendations (Bennett et al. 2016; Roni et al. 2013). From extensive surveys here and in Polivka et al. (2015), it is clear that detection of reach-scale capacity increase vs. basic density increase requires: 1) separate surveys of restored and unrestored habitat *within* a treated reach and 2) availability of and access to physically and biologically similar untreated reaches. The first stipulation suggests that reach scale surveys might not be effective at detecting Type 1 capacity increases if unrestored habitat within the treated reach obfuscates the effect of restoration, or when the effect of ELJs is very localized (Polivka et al. 2019). The second stipulation means that the characteristics of control reaches should thus be a consideration both for restoration site selection and for post-restoration effectiveness monitoring designs.

The inability to distinguish between redistribution and capacity increase following restoration due do study design or scale issues is a possible explanation for inconsistent responses (Smokorowski and Pratt 2007; Whiteway et al. 2010), small or no observed effects of restoration (Roni et al. 2008; Stranko et al. 2012; Hillman et al. 2016), or positive responses that vary by species, year, and time period within a rearing season (Pess et al. 2012; Polivka et al. 2015). We have shown that juvenile salmonid habitat capacity increased via the Type 1 outcome (Figure 1), distinguishable from simple redistribution. Studies that are more mechanistic in nature are required to support observations of capacity increases here and in related studies (Beechie et al. 2015), such as linking capacity increases to improvements in traits correlated with fitness, such as sub-yearling growth or survival through any subsequent life stage (Achord et al. 2007; Polivka et al. *in review*). Restoration in spawning streams is implemented to augment habitat for sub-yearling, an important stage of the “stream-type” life history when considerable growth and development take place in streams (Groot and Margolis 1991). The apparent capacity increase described here for the upper and lower geomorphic basin segments of the Entiat River indicate that restoration is benefitting Chinook salmon and, to a less definite extent, steelhead early in their life history.

## [A]Acknowledgements

Funding for this work was provided by the US Bureau of Reclamation. Cascadia Conservation District (CCD) provided field assistants to support this work via a series of Joint Venture Agreements with PNW Research Station. We thank P. Entzel and V. Hampton at CCD for administrative support for those agreements. J. Novak, H. Porter and R. Hosman led the collection of field data with assistance from S. Eichler, R. Logan, K. Logan, J. Jorgensen, H. Porter, B. Forney, S. Letzing, R. Hosman, K. Swieca, O. Graham, N. Holt, S. Kaech, and C. Skalisky. Assistance with Bayesian analysis was provided by G. Stewart. Helpful comments on drafts of this manuscript were provided by P. Roni, R. Flitcroft, T. Hillman and two anonymous reviewers.

